# Nitric oxide donor sodium nitroprusside serves as a source of iron supporting *Pseudomonas aeruginosa* growth and biofilm formation

**DOI:** 10.1101/2025.03.12.642859

**Authors:** Xavier Bertran i Forga, Yaoqin Hong, Kathryn E. Fairfull-Smith, Jilong Qin, Makrina Totsika

## Abstract

Biofilm dispersal agents, like nitric oxide (NO), restore antimicrobial effectiveness against biofilm infections by inducing bacteria to shift from a biofilm to a planktonic state, thereby overcoming the antimicrobial tolerance typically associated with biofilms. Sodium nitroprusside (SNP) is a widely used NO donor for investigating the molecular mechanisms underlying NO-mediated biofilm dispersal in the nosocomial pathogen *Pseudomonas aeruginosa*. However, the biofilm effects of SNP are variable depending on the *in vitro* experimental conditions, with some studies reporting enhanced growth in both planktonic and biofilm forms instead of dispersal. These discrepancies suggest that SNP affects *P. aeruginosa* biofilm-residing cells beyond the release of NO. In this study, we compared SNP with another NO donor, Spermine NONOate, to systematically contrast their effects on biofilm and planktonic cultures of *P. aeruginosa*. We found that SNP, but not Spermine NONOate, increased the biomass of *P. aeruginosa* biofilms in microplate cultures. This effect was also observed when biofilms were supplemented with iron. Additionally, supplementation with SNP rescued the planktonic growth of *P. aeruginosa* in iron-depleted media, similar to FeSO_4_ supplementation, suggesting that SNP may serve as an iron source. Our findings suggest that SNP’s potential as an NO agent used for biofilm dispersal may be confounded by its role in promoting both biofilm and planktonic growth through its iron centre. Our study cautions investigators using SNP for studying NO-mediated biofilm dispersal.

## Importance

Research into biofilm dispersal agent nitric oxide (NO) holds promise for treating biofilm-associated infections. Sodium nitroprusside (SNP), an NO donor widely used in antibiofilm research, has been shown in this study to enhance cell growth and biofilm formation in *P. aeruginosa* by acting as a source of iron. Our results suggest that SNP functions both as NO and iron donor, with its iron-releasing properties playing a more dominant role in promoting biofilm growth in closed culture systems. This study underscores the dual but conflicting roles of SNP in biofilm growth, which caution its future development as a NO-based therapeutic strategy for biofilm-associated infections.

## Observation

Biofilms are microbial communities encapsulated in an extracellular polymeric matrix that are ubiquitous in both natural and clinical environments. They cause significant damage through biofouling of equipment and serve as a reservoir for recurrent chronic infections, as well as for food and water contamination^1^. In addition, bacteria in biofilms exhibit increased tolerance to antimicrobials and disinfectants compared to their planktonic counterparts, which undermines therapeutic efficacy^2^. One approach to address the substantial economic and health challenges posed by bacterial biofilms is to induce a transition to the planktonic state using biofilm dispersal agents, thereby reducing their antimicrobial resistance. Nitric oxide (NO), a well-known biofilm dispersal agent, has been exploited as an antibiofilm agent against several clinically and industrially relevant biofilm-forming species ^3,4^. In *Pseudomonas aeruginosa*, NO functions as a signalling molecule, promoting the degradation of the exopolysaccharide matrix and the assembly of flagella and therefore reverting biofilm cells to the planktonic (free-living) state^5,6^. As a result, NO has been shown to enhance the effectiveness of antibiotics by resensitising biofilm bacteria to antimicrobial treatment, aiding in biofilm eradication through this shift to the planktonic form ^4,7^. In *P. aeruginosa, Escherichia coli* and *Staphylococcus aureus*, NO has been reported to disrupt Fe-S clusters through S-nitrosylation or heme-containing proteins, which are implicated in cell signalling and metabolic processes ^8–11^. The reaction of oxygen radicals with NO leads to the formation of peroxynitrite, a highly reactive nitrosative species that may damage lipids and DNA ^12^.

The precise delivery of gaseous NO at desired concentrations has remained a challenge in both clinical and laboratory settings due to its gaseous and highly reactive nature (14). This has prompted the development of NO-releasing drugs, which are generally soluble compounds that have aided in the controlled delivery of NO and have thus been instrumental to investigate the mechanisms involved in associated biofilm dispersal responses. The metal-nitrosyl complex sodium nitroprusside (SNP), an FDA-approved vasodilator, has been a model compound in studying NO-induced biofilm dispersal mechanisms in *P. aeruginosa*^5–7,13,14^. SNP consists of a ferrous (Fe^2+^) ion coordinating five cyanide groups and a nitrosonium group (NO^+^), which is released as NO together with cyanide upon reduction in aqueous solution (Fig 1A) ^15^. Due to its reported success in dispersing biofilms *in vitro*, SNP is currently being explored in combination with nanoparticles as an *in-situ* NO-delivery method with antibiofilm properties using a mouse skin infection model ^16^. However, biofilms grown in closed culture systems (i.e., microtiter plates) showed confounding results following 12-24h of SNP treatment, with effects ranging from growth promotion of planktonic and biofilm bacteria to biofilm biomass reduction^17–19^. In contrast, other NO donors, such as *N-* diazenium diolates (NONOates), are consistently reported to reduce the biomass of biofilms *in vitro* ^11,17^. A primary distinction between these two NO donors is the ferrous iron present in SNP that is lacking from NONOates. Iron has been consistently reported to aid in the formation of *P. aeruginosa* biofilms^20,21^. Therefore, we hypothesised that the variable effects reported for SNP on biofilms *in vitro* may be substantially influenced by the other components in its chemical structure.

**Fig 1.**
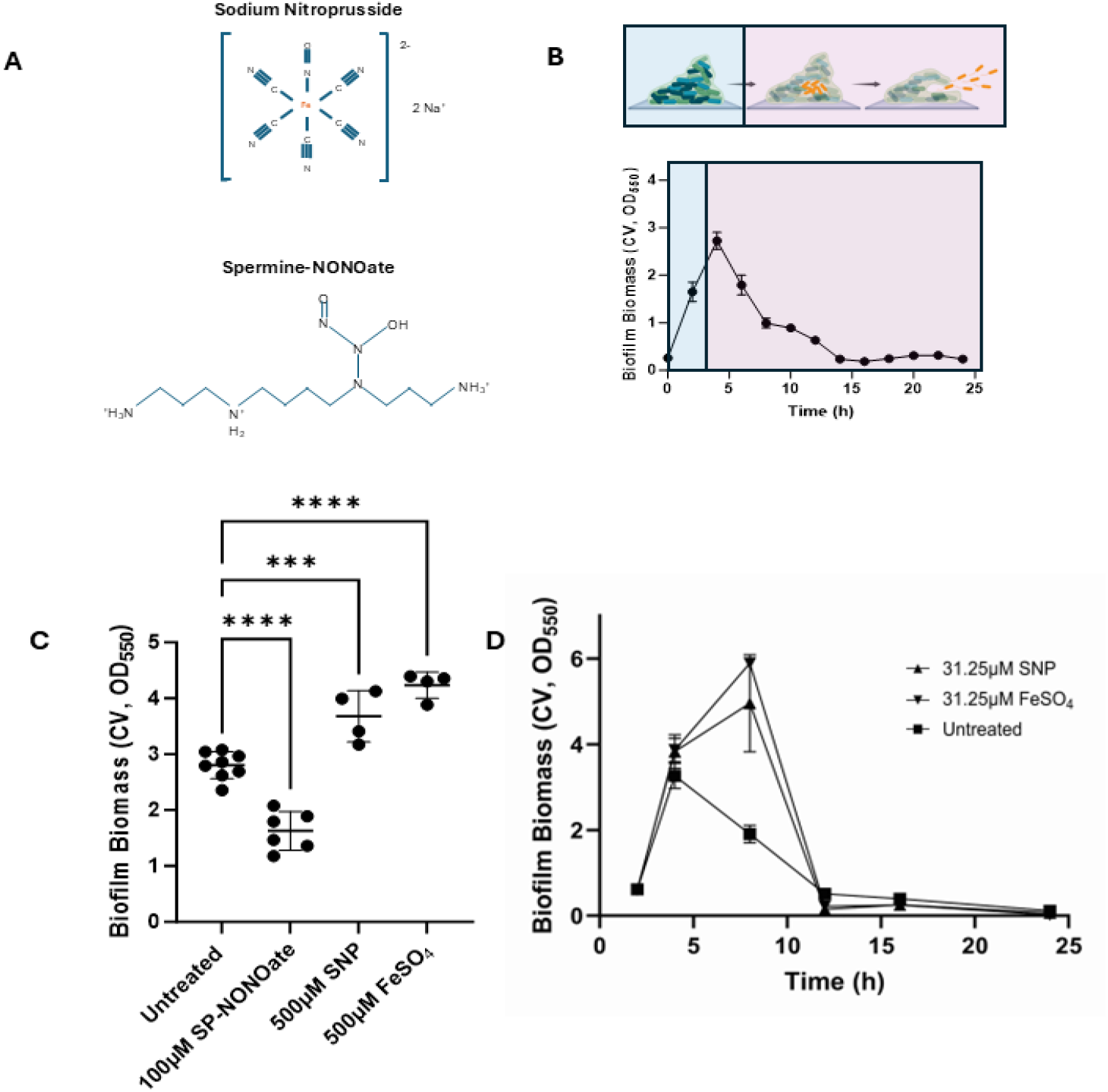
SNP promotes biomass increase of *P. aeruginosa* PAO1 biofilms. **(A)** Chemical structure of Sodium Nitroprusside and Spermine-NONOate. **(B)** Changes in biofilm biomass over 24h. 10^7^ CFU/ml bacterial solutions were prepared from overnight cultures using M9 media, transferred to 24-well plates and incubated at 37 ºC shaking. Biofilm biomass was quantified every 2h by crystal violet staining for up to 24h. **(C)** 4h biofilms were treated with SP-NONOate for 15-min, or FeSO_4_ or SNP for 30-min, at the indicated concentrations. Data represent at least four independent replicates. **(D)** 4h biofilms of either PAO1 WT or *fhp*::Tn were treated with either SP-NONOate or SNP for 15-min or 30-min respectively at the indicated concentrations. Data represent two independent replicates. **(E)** Biofilm cultures were grown in M9 media supplemented with 31.25µM of SNP or FeSO_4_. Two independent cultures were included. The means ± SD are represented in the graph.

### *SNP increases biofilm biomass of P. aeruginosa PAO1* in vitro

To explore the effects of SNP on microplate biofilms of *P. aeruginosa* (a model biofilm-forming pathogen), we first measured biofilm growth kinetics in minimal media by tracking the biomass of *P. aeruginosa* PAO1 biofilms over 24 hours in microtiter plates (Fig 1A, Text S1). In this system, *P. aeruginosa* biofilms reached a peak biomass at 4 hours (biomass_max_), followed by a gradual decline, with <10% biofilm biomass remaining by 14 hours. As the biomass_max_ achieved in microplates occurs at 4 h of incubation, 4h-old biofilms were subsequently used to evaluate dispersal effects by two NO-donors, SNP and Spermine NONOate (SP-NONOate), measured as significant reduction in remaining biofilm biomass.

SP-NONOate is an *N-*diazeniumdiolate that spontaneously releases NO in aqueous solution ^11,17,22^. Therefore, SP-NONOate was used to compare the effects of SNP on biofilm biomass. Consistent with previous reports, SP-NONOate induced a reduction of *P. aeruginosa* biofilm biomass within 15 minutes (Fig 1C, Fig S1A and S1B) ^22^. Instead, SNP induced a dose-dependent biofilm biomass increase after 30 minutes, with a maximum 1.25-fold increase observed at 500 µM SNP (Fig 1C, Fig S1B).

As a treatment strategy, NO is proposed to reduce biofilm-associated antimicrobial tolerance by reverting *P. aeruginosa* biofilm cells to their planktonic state. This hypothesis has been routinely tested using SNP to induce NO-mediated dispersal of *P aeruginosa* biofilms ^5,6,14,17^. However, our findings contradict this expectation, as we observed that SNP instead promotes a rapid increase in biofilm biomass. A previous study similarly reported that short exposure to low concentrations of NO correlated with increased *P. aeruginosa* biofilm biomass ^11^, suggesting that SNP might be releasing NO at concentrations insufficient for dispersal, thereby promoting biofilm growth instead. To investigate this possibility, we treated PAO1 biofilms with a range of sub-dispersing concentrations of SP-NONOate (Fig. S2), yet observed no significant biomass increase. These results suggest that SNP may enhance biofilm growth through a mechanism independent of NO release.

A major difference in chemical structures between SNP and SP-NONOate is the presence of an iron centre in SNP (Fig 1A). Iron has been described as an essential micronutrient for the development of biofilms in *P. aeruginosa* ^20^. Therefore, we reasoned that the observed increase in biomass by SNP (Fig 1C) could be derived from the surplus in bioavailable iron released from the SNP molecule. This hypothesis is supported by previous reports showing that *P. aeruginosa* biofilms grown in iron-supplemented media reached a higher biomass, and that the addition of iron to established biofilms induced rapid surface attachment of planktonic cells ^21,22^. Therefore, we supplemented established *P. aeruginosa* biofilms with matching concentrations of FeSO_4_, which induced a similar increase in biofilm biomass to that of SNP treatment (Fig 1C, Fig S1B). Unlike SP-NONOate, which significantly reduced biofilm biomass accumulation (Fig S3), media supplementation with SNP or FeSO_4_ prolonged the biofilm formation phase to 8 h compared to the control media group (4 h), allowing for increased accumulation of biomass and a higher biomass_max_ (Fig 1E). Therefore, these data suggest SNP as a potential iron donor to support biofilm growth.

### SNP rescues growth in iron-depleted conditions

To further test this hypothesis, we next examined whether supplementing iron-chelated M9 medium with SNP could rescue the growth of *P. aeruginosa* planktonic cultures. While M9 supported the growth of *P. aeruginosa*, we found that growth in M9 was inhibited when iron was depleted from culture media by the iron chelator 2,2’-bipyridyl (Fig 2, Text S1). However, the addition of SNP reversed the growth inhibition caused by iron depletion, and that this rescue of otherwise defective growth of PAO1 in iron-chelated media, by the addition of SNP, was comparable to the effect of FeSO_4_ supplementation in alleviating growth arrest triggered by iron deficiency (Fig 2, Fig S4). Together, these data suggest that SNP acts as an iron donor, supporting both biofilm and planktonic growth of *P. aeruginosa*.

**Fig 2.**
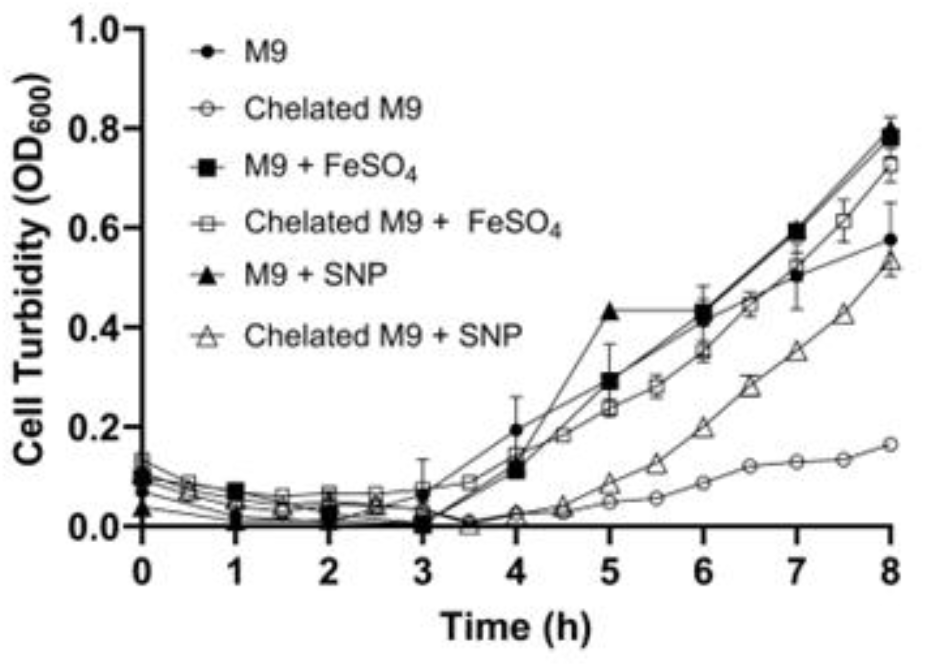
Growth defects of *P. aeruginosa* PAO1 in iron-depleted media can be rescued by SNP or FeSO_4_. Initial bacterial cultures were inoculated at an OD_600_ of 0.05 and grown in M9 media pre-treated with 100µM of the iron scavenger 2,2’-bipyridyl. Cultures were supplemented with SNP or FeSO_4_ under shaking conditions in 96-well microtiter plates. Culture optical density was periodically recorded in a plate reader by measuring the absorbance at 600nm. Data represents two biological replicates. The means ± SD are represented in the graph.

While the mechanism by which SNP releases iron remains unclear and is beyond the scope of this work, our data suggest that physiologically relevant iron concentrations become readily available upon SNP addition, potentially overshadowing its NO-related effects. This is evidenced by the rapid increase in biofilm biomass within 30 minutes of SNP exposure, similar to that induced by FeSO_4_, possibly due to the upregulation of extracellular matrix polysaccharide Psl synthesis and/or enhanced attachment of planktonic bacteria ^22^. Importantly, our findings highlight the iron-releasing properties of SNP, a factor that may not have been fully considered in previous studies investigating its therapeutic potential, likely introducing bias to our understanding of SNP as a NO-donor. While studies contrasting the phenotypic and genotypic behaviours of biofilms grown in each culture system are lacking, existing data suggest that biofilms grown in open flow systems exhibit distinct phenotypic profiles compared to those in closed microplate environments. This is evident from the opposing responses of *P. aeruginosa* biofilms to iron chelators and tobramycin, depending on the culture system used^23^. These findings highlight the value of microplates as a low-cost, rapid, and high-throughput system usable to control for potentially masked effects by treatments tested in open cultures.

In terms of potentiating antibiofilm treatments, SNP was reported previously to synergise with tobramycin against *P. aeruginosa* biofilms, an antibiotic commonly used to treat biofilm infections in cystic fibrosis patients^7,24^ and more effective in metabolically active bacteria^27,28^. Here we showed that SNP supplementation resulted in a higher growth rate for *P. aeruginosa* due to its iron donation, which could lead to a more active metabolism, potentially resulting in sensitisation of biofilm-embedded bacteria to tobramycin. In addition, iron supplementation has also been shown to increase the effectiveness of antibiotics by enhancing the production of reactive oxygen species. This would also explain the reported increases in effectiveness of nanoparticles when tethered to SNP, as these depend on light-activated ROS generation^25^. Considering that SNP acts as a source of readily available iron, ROS-induced damage to the biofilms might have been enhanced by the presence of ferrous iron in SNP, which may trigger localised production of superoxide radicals through the Fenton reaction leading to a severe disruption of iron homeostasis^25^.

## Conclusion

Altogether, our results demonstrate that SNP provides a readily available source of iron to growing biofilms of *P. aeruginosa*. Considering most assays studying NO dispersal have been conducted with SNP as the NO donor, our findings should elicit caution when solely attributing reported SNP biofilm responses to NO, as masked secondary effects might have been overlooked.

## Acknowledgements

This work is funded in part by an Australian Research Council project grant (DP210101317), the Max Planck Queensland Centre on the Materials Science of Extracellular Matrices to MT, and the QUT Amplify Scholarship provided by the Queensland University of Technology (Australia) to XB. The Ian Potter Foundation sponsored the CLARIOStar high-performance microplate reader (BMG, Australia). The funders had no role in study design, data collection and analysis, decision to publish, or preparation of the manuscript. The authors would like to thank Professor Robert EW Hancock (University of Columbia) for providing the *P. aeruginosa* PAO1 strain used in this study.

## Author contributions

XB and JQ conceptualised the project. XB, JQ and YH contributed to experimental design. XB conducted all experiments, and contributed to data collection, analysis and visualisation. XB, JQ, YH and MT contributed to data interpretation. JQ and MT supervised the project. KFS and MT obtained the funding. XB wrote the original draft, and all authors edited the manuscript.

## Data availability

All data generated or analysed during this study were included in this article and supplementary files.

